# Extracellular Vesicle Biomarkers in the Aqueous Humor Correlate with Retinoblastoma Seeding

**DOI:** 10.64898/2026.06.01.728823

**Authors:** Chen-Ching Peng, Anne Amacker, Nerea Goni, Jyothi Attem, Shreya Sirivolu, Jesse L Berry, Liya Xu

## Abstract

**Purpose:** To investigate the correlation of retinoblastoma (RB) aqueous humor (AH)–derived extracellular vesicle (EV) surface markers, particularly CD133, with clinical features of RB.

**Methods:** AH samples (n=22) collected at diagnosis (n=17) or primary enucleation (n=5) from RB eyes were profiled using magnetic bead-based flow cytometry. Median fluorescence intensities (MFI) of tetraspanins (CD9, CD63, CD81) and CD133 were quantified and correlated with International Intraocular RB Classification (IIRC) stage, seeding location, seeding morphology, and enucleation.

**Results:** CD133^+^ EVs were detected in 16 of 22 (73%) samples. While CD133 levels showed a non-significant increasing trend with advanced IIRC stages, CD133 expression was significantly elevated in eyes with vitreous seeding vs. subretinal only (P = 0.010). Cuzick’s trend analysis revealed a significant association between rising CD133 % MFI and complex seeding (P < 0.001). Conversely, CD63 displayed a significant inverse trend, with decreasing % MFI associated with complex seeding (P < 0.001). CD133^high^ eyes (MFI>10,000, n=9) exhibited increased enucleation (P = 0.007) and predominantly sphere/cloud seeding patterns (P = 0.001). CD133^high^ eyes demonstrated a coordinated CD63/CD81 co-expression pattern (r = 0.80), absent in the CD133^low^ group.

**Conclusions:** Elevated CD133 level and percentage were associated with vitreous involvement and complex seeding morphology, suggesting potential as a minimally invasive indicator of disease burden. The inverse relationship between CD63 and seeding severity, combined with CD63/CD81 remodeling in CD133^high^ eyes, suggests an EV profile indicative of aggressive disease. These findings support the association of CD133^+^ EVs with advanced RB seeding and warrant further prospective validation.

## Introduction

Retinoblastoma (RB) is the most common pediatric intraocular malignancy^1^ accounting for about 3% of all childhood cancers^2^. Around 300 cases in the United States and 8,000 global cases are diagnosed every year^2,3^. The goal of treatment is to save the child’s life and if possible, save the eye with some degree of vision. Traditional tumor biopsy in RB is contraindicated due to the risks of extraocular spread of the cancer^4-9^; thus, tumor tissue is available only at enucleation. This limits understanding of the disease at both the genotypic and phenotypic level^4-9^ and hinders application of genomics to treatment decision making due to lack of in vivo RB markers from eyes undergoing therapy^10^. These challenges highlight the clinical need for innovative methodologies.

A significant breakthrough in 2017 demonstrated that aqueous humor (AH), an ocular fluid in direct contact with intraocular tumors, could serve as a source of tumor-derived cell-free DNA (cfDNA), enabling safe, minimally invasive liquid biopsy in RB. However, limitations to analysis due to factors such as decreasing DNA concentrations with ongoing treatment^11,12^ highlight the need for complementary biomarkers that reflect tumor biology through mechanisms beyond genomic alterations. Beyond cfDNA, extracellular vesicles (EVs) have emerged as important mediators of intercellular communication and hold promise as non-genomic biomarkers^13,14^. EVs are heterogeneous nanosized lipid membrane-enclosed vesicles secreted by all kinds of cells into biological fluids, carrying proteins, nucleic acids, and lipids. EVs are highly stable due to their protection by a lipid bilayer. They reflect the status of the cell, rendering them suitable as a biomarker^15^, with presence of integral membrane protein markers like tetraspanins (e.g., CD9, CD63, and CD81) serving as common EV markers due to their functional role and extensive presence. These vesicles are abundant and found in various ocular fluids, including the AH^16,17^, making them accessible through minimally invasive sampling.

Our previous work established that AH in a subset of RB eyes contains a distinct population of CD133-enriched EVs, a known cancer stem cell marker^18^, not present in glaucoma or uveal melanoma samples, supporting their tumor specificity^19^. This was validated using multiple EV characterization platforms, including MACSPlex, Single-particle interferometric reflectance imaging sensor (SP-IRIS), and Single Extracellular VEsicle Nanoscopy (SEVEN) assays^19^. MACSPlex is a multidimensional bead-based assay with 39 antigens and two controls tied to APC conjugated CD9, CD63, and CD81 antibodies, allowing extensive investigation of known cancer and cell type markers. We found that CD133 is frequently co-localized with CD63 and CD81, supporting a unique RB-derived EV signature^17^. While tetraspanins serve as useful pan-EV surface markers, they may not provide the precision needed to discern RB-derived EVs (RB-EVs) from other diseases or differentiate disease subtypes within RB. Thus, tumor-specific surface markers such as CD133 may offer enhanced specificity in profiling RB disease state. However, CD133 expression across RB samples was heterogeneous, and one subset of cases demonstrated little to no detectable CD133. This variability raised the question of whether CD133^+^ EVs reflect clinically meaningful features of disease behavior.

Tumor seeding, defined as the spread of RB cells into adjacent liquid or semi-liquid spaces, is a key determinant of prognosis and treatment complexity^20^. This seeding has been described as both vitreous and subretinal based on the anatomical location into either the vitreous chamber or behind the retina. Seeding also can be classified by breadth of involvement, using a quadrant system of 0-4 to describe involvement. Seeding plays a central role in RB staging, with its presence categorizing eyes as group C or higher in the International Classification for Intraocular RB (IIRC)^21^. Clinically, vitreous seeds are particularly challenging to treat due to their isolation from systemic circulation, limiting chemotherapy efficacy^22^. Both vitreous and subretinal seeding are major contributors to treatment failure and enucleation, even with advances in localized therapies^23-25^. Given the biological importance of seeding morphology and extent, an AH-based biomarker capable of reflecting these features could provide clinically actionable information.

Building on our initial observations, we hypothesized that CD133 signal intensity on AH-derived EVs correlates with clinically relevant features of RB including seeding location, morphology, and enucleation. In this study, we analyzed a diagnostic cohort using multiplexed bead-based EV profiling (MACSPlex) to determine whether CD133^+^ EVs correlate with complex tumor seeding.

## Methods

### Patients and Sample Collection

This study was conducted under the IRB approval at USC and Children’s Hospital Los Angeles (CHLA). CHLA maintains the following active CHLA IRB approval: CHLA-17-00248, RB Patient Clinical Database, and Tissue Biorepository and all procedures adhered to institutional and federal guidelines. The methods used in this study adhered to the tenets of the Declaration of Helsinki.

All patients included in the analysis provided written informed consent for the biorepository at CHLA via a legal guardian. AH data was kept separate from clinical data until the final retrospective analysis. AH samples were collected via clear corneal paracentesis from 22 RB eyes at time of diagnosis (n=17) or primary enucleation (n=5) from 19 patients. Clinical data was masked from the laboratory team until after EV analysis was completed. All AH samples were collected and stored at −80 °C for analysis. No pre-enrichment or sample cleanup was required prior to MACSPlex analysis. A fixed input volume of 20 μL was used for all samples to minimize variability in signal intensity.

### Broad-Based Multiplex Flow Cytometry Assay (MACSPlex)

AH samples were processed and analyzed following the general principles outlined in the Minimal Information for Studies of Extracellular Vesicles (MISEV2023) guidelines^26^, although no additional EV enrichment steps were performed. Briefly, AH samples were first centrifuged to remove cellular debris, and the supernatant was directly subjected to bead-based multiplex analysis by flow cytometry (MACSPlex Exosome Kit, human, Miltenyi, Gaithersburg, MD, USA) per the manufacturer’s instructions and as described in our previous publication^19^. Up to 20 μL of AH was added to a 1.5 mL tube. MACSPlex Buffer Solution was added to each tube until the total volume reached 120 μL. MACSPlex Exosome Capture Beads (CD1c, CD2, CD3, CD4, CD8, CD9, CD11c, CD14, CD19, CD20, CD24, CD25, CD29, CD31, CD40, CD41b, CD42a, CD44, CD45, CD49e, CD56, CD62P, CD63, CD69, CD81, CD86, CD105, CD133, CD142, CD146, CD209, CD326, HLA-ABC, HLA-DRDPDQ, MCSP, ROR1, and SSEA-4) were vortexed for 30 seconds, and 15 μL was added to each tube. Tubes were then incubated overnight at room temperature (RT) in an orbital shaker at 450 rpm while protected from light. After incubation, 500 μL of MACSPlex Buffer was added to each tube and centrifuged at RT for 5 min at 3000×g. Following centrifugation, 500 μL of supernatant was removed, and 15 μL of the fluorescent antibody cocktail (CD9, CD63, and CD81) was added and mixed by pipetting. Samples were then incubated for 1 hour under the same conditions. Subsequently, buffer wash, and centrifugation steps were repeated twice as described, and samples were finally resuspended in 400 μL of MACSPlex Buffer for flow cytometry analysis using the FACSymphony S6 cell sorter (Becton Dickinson, Franklin Lakes, NJ, USA) at the CHLA TSRI Flow Cytometry Core. A workflow graphic is available in our previous publication^19^.

The median fluorescence intensity (MFI) values of each marker were corrected for background signal by subtracting the respective MFI values from the non-EV-containing buffer samples included in every session of analysis. Thus, no signal was defined as a value that was less than or equal to the background signal from control buffers. All samples were then normalized to 20 μL for comparison of fluorescent signal intensities.

### Statistical Analysis

Continuous variables (MFI or marker percentages) were summarized as mean ± standard error of the mean (S.E.M.). Data normality was assessed using the Shapiro–Wilk test, and non-parametric methods were applied throughout for non-normally distributed variables. For comparisons involving more than two groups (Figures 2 and 3), overall significance was determined using the Kruskal–Wallis (K–W) test, followed by Dunn’s multiple-comparison post-hoc analysis. Ordered categorical variables, seeding morphology (none, subretinal only, dust, sphere, and cloud), were evaluated using Cuzick’s non-parametric trend test to assess monotonic changes across increasing seeding complexity (Figure 4 and Supplementary Figure 1). Two-group comparisons, including enucleation status and seeding types between CD133^low^ and CD133^high^ subgroups, were analyzed using the Fisher’s exact test and the Cochran–Armitage test, respectively (Figure 5B). Correlative analyses among tetraspanin markers (CD63, CD81) were performed in CD133^low^ and CD133^high^ samples using Pearson’s correlation coefficients (Figures 5C) and linear regression models were fitted to evaluate concordance between CD63% and CD81% expression levels. All statistical tests were two-tailed, and P < 0.05 was considered statistically significant. Analyses were performed using GraphPad Prism version 10.0 (GraphPad Software, Boston, MA, USA) and R version 4.3.3 (R Foundation for Statistical Computing, Vienna, Austria).

## Results

### Demographic and clinical features of the diagnostic cohort of RB patients

To correlate clinical and molecular data, we compiled demographic and disease-specific features from each eye. These features included RB1 germline mutation status, laterality, age at diagnosis, sex, tumor growth pattern, presence of subretinal or vitreous seeding, and enucleation status. AH samples were grouped by IIRC stage at diagnosis and further stratified by seeding extent, defined as the number of quadrants involved (0–4; Figure 1A). Each AH sample was analyzed independently. Two patients contributed bilateral samples, which were treated as separate data points for EV and clinical comparisons (Figure 1A). Using MACSPlex analysis, a heatmap of the 22 samples demonstrated robust detection across all antigens, with notable co-dominance of CD63 and CD81. CD133 expression exhibited substantial variability, with 6 samples negative for CD133 and 16 samples positive for CD133, showing varying signal intensities (Figure 1B). A representative MACSPlex profile is shown in Figure 1C.

**Figure 1.**
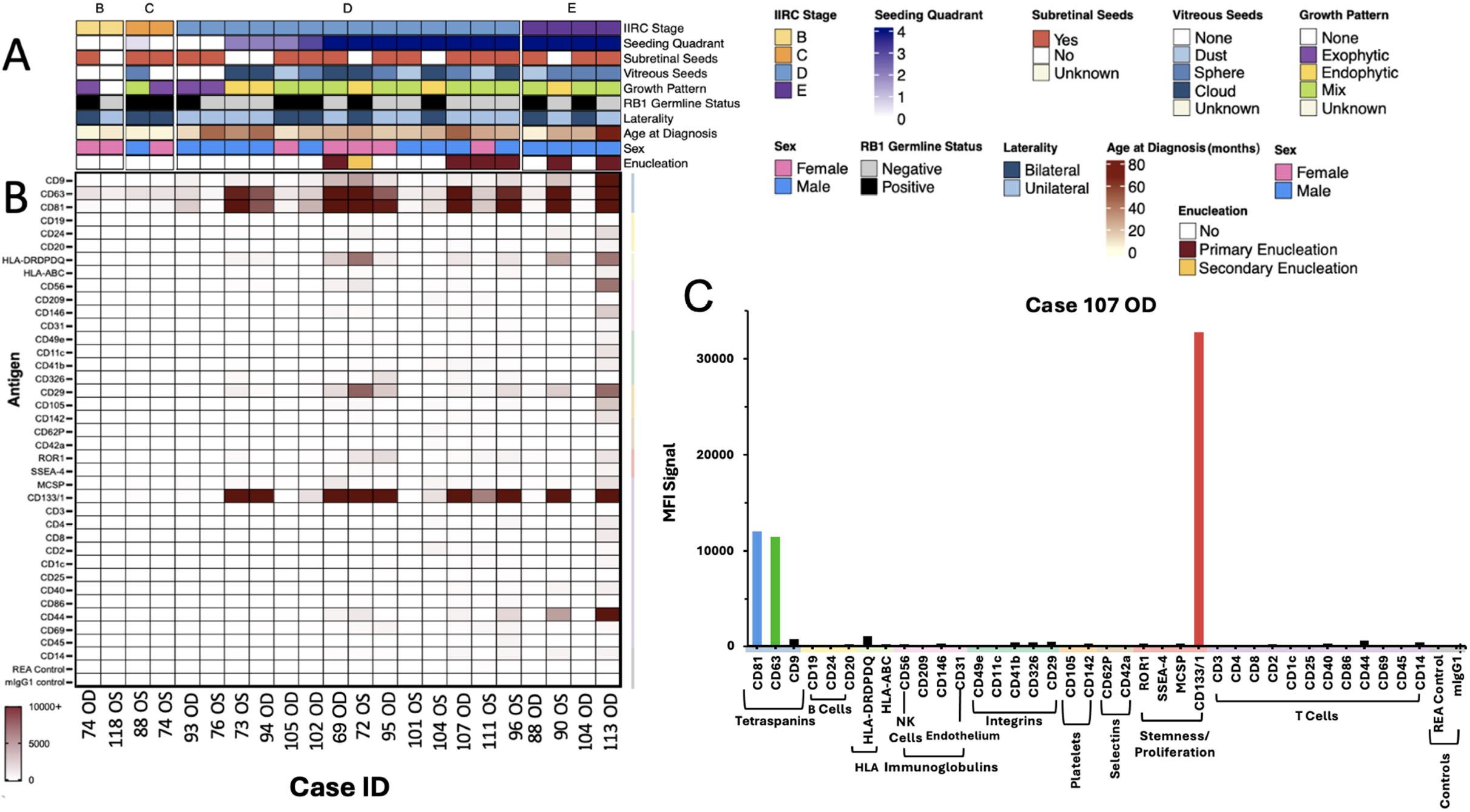
Demographic and clinical variables for the RB AH cohort and MACSPlex analysis. (A) Twenty-two AH samples (5 from primary enucleation and 17 at diagnosis) were grouped by IIRC stage, with various clinical variables color-coded, including RB1 germline mutation status, laterality, age at diagnosis, sex, growth pattern, presence of vitreous or subretinal seeding, seeding quadrant, and enucleation status. (B) An MFI heatmap was created for all samples, displaying the expression levels of 39 EV-associated antigens. MFI signals were normalized to an input volume of 20 µL and visualized using a color scale ranging from 0 to 10,000 MFI, with values above 10,000 displayed at the maximum intensity of the scale (C) An example of a MACSPlex profile generated from Case ID 107 OD.

### Tetraspanin and CD133 expressions relative to IIRC stage

To assess whether EV marker expression correlates with tumor burden, we analyzed the levels of CD63, CD81, CD9, and CD133 in AH-derived EVs from RB patients stratified by IIRC stage (Figure 2). While CD63 expression (Figure 2A) appeared elevated in several stage E samples compared with earlier stages, Kruskal–Wallis analysis demonstrated no statistically significant differences among groups (P = 0.320). Similarly, CD81 expression (Figure 2B) showed a progressive increase across disease stages, and overall group comparison reached statistical significance (P = 0.029). CD133 expression (Figure 2C) also trended higher in stage E samples, although this did not reach statistical significance (P = 0.059). Likewise, CD9 expression (Figure 2D) increased with advancing stage and demonstrated a significant overall Kruskal–Wallis result (P = 0.016). However, despite significant overall group differences for CD81 and CD9, Dunn’s post hoc pairwise comparisons did not identify statistically significant differences between any individual stage pairs for any marker. Although tetraspanin and CD133 expressions across stages did not show statistical differences, the observed trends were biologically consistent. This prompted further exploration of tetraspanin and CD133 signals in relation to clinically important features, particularly focusing on tumor seeding.

**Figure 2.**
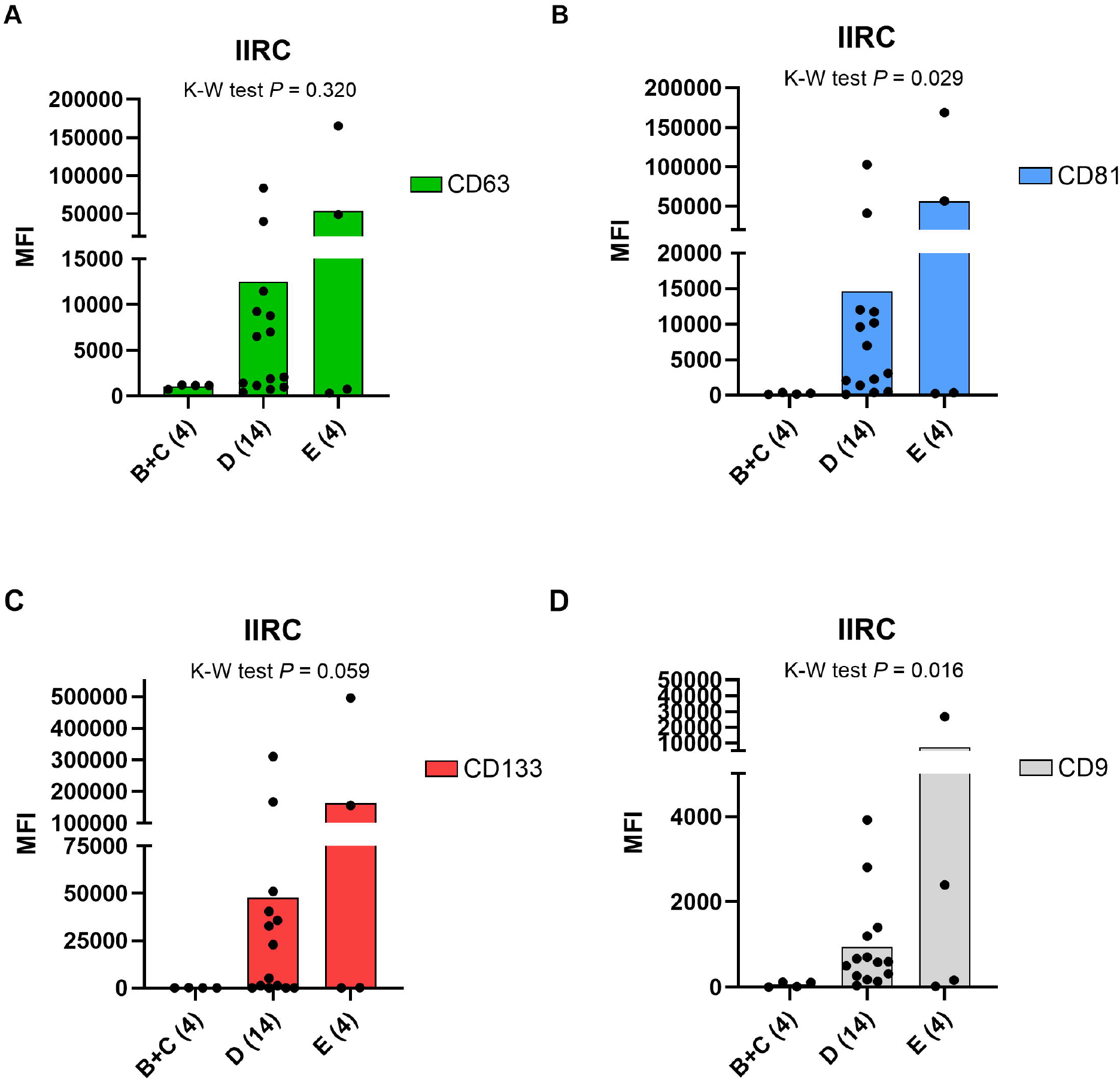
Tetraspanins and CD133 expression levels in AH-derived EVs stratified by IIRC stage. The MFI of (A) CD63 (green), (B) CD81 (blue), (C) CD133 (red), and (D) CD9 (grey) was quantified in AH-derived EVs from RB patients, categorized by International Intraocular RB Classification (IIRC) stage: B (n = 2), C (n = 2), D (n = 14), and E (n = 4). Statistical differences across IIRC stages for CD63, CD81, CD133, and CD9 were assessed using Kruskal-Wallis (K-W) tests, with pairwise comparisons conducted using Dunn’s multiple comparisons test. Data are presented as mean ± SEM, with individual samples represented as dots.

### EV markers in relation to seeding location

To determine whether AH-derived EV markers reflect intraocular tumor dissemination patterns, we examined CD63, CD81, CD9, and CD133 expressions in samples grouped by seeding location (Figure 3A): subretinal only (n = 4), vitreous only (n = 5), or both subretinal and vitreous (n = 12), with one case with no seeding. CD63 expression demonstrated a trend toward increased levels in eyes with vitreous seeding compared to those with subretinal only disease (Figure 3B), although the differences were not statistically significant (P = 0.083). A similar pattern was observed for CD81 (Figure 3C), with the highest levels seen in eyes with both subretinal and vitreous involvement (P = 0.050), although post hoc pairwise comparisons did not reach significance. In contrast, CD133 expression was significantly elevated in eyes with vitreous seeding compared to subretinal only cases (P = 0.006; Figure 3D). Pairwise analysis confirmed significantly higher CD133 levels in the vitreous only group compared to subretinal only (P = 0.010), suggesting that CD133 may serve as a specific EV biomarker associated with vitreous dissemination in RB. Statistical significance was not present for CD9 (Figure 3E), with highest levels in samples containing both seeding types (P = 0.140). In the one case without seeding, MFI values for all markers were included (Figure 3F). These results indicate that among the EV markers analyzed, CD133 most strongly reflects vitreous tumor seeding and may serve as a liquid biopsy marker for identifying advanced intraocular spread.

**Figure 3.**
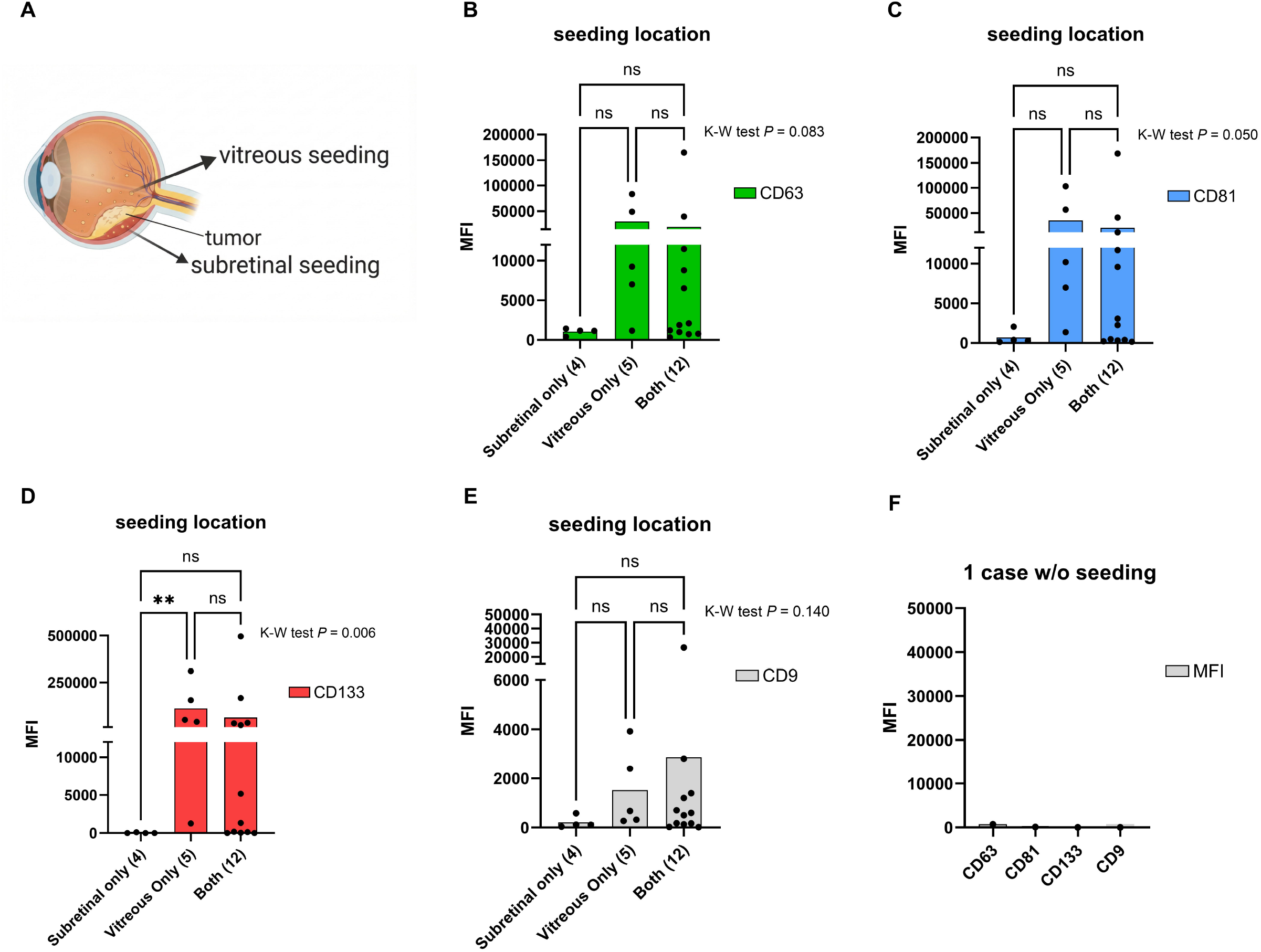
Tetraspanins and CD133 expression levels in AH-derived EVs stratified by seeding location. (A) A schematic illustrating intraocular tumor dissemination patterns, including subretinal and vitreous seeding. (B–E) MFI of CD63 (green, B), CD81 (blue, C), CD133 (red, D), and CD9 (grey, E) in AH EVs from RB eyes categorized by seeding location: subretinal only (n = 4), vitreous only (n = 5), or both sites (n = 12). Kruskal-Wallis (K-W) tests assessed statistical differences across locations for CD63, CD81, CD133, and CD9, with pairwise comparisons conducted using Dunn’s multiple comparisons test. Data are presented as mean ± SEM, with individual samples represented as dots. “ns” indicates not significant. (F) CD63, CD81, CD133, CD9 MFI signal intensity for the 1 case included with no seeding present.

### EV markers and seeding morphology

To further evaluate the relationship between vitreous seeding characteristics and EV biomarker levels, we stratified AH samples by dominant seeding morphology: none (n = 1), subretinal only (n = 4), dust (n = 4), sphere (n = 7), and cloud (n = 6). Seeding morphologies were illustrated in Figure 4A. Signals were converted to percentages (marker MFI over total MFI), thereby reducing bias introduced by differences in total EV content across samples.

**Figure 4.**
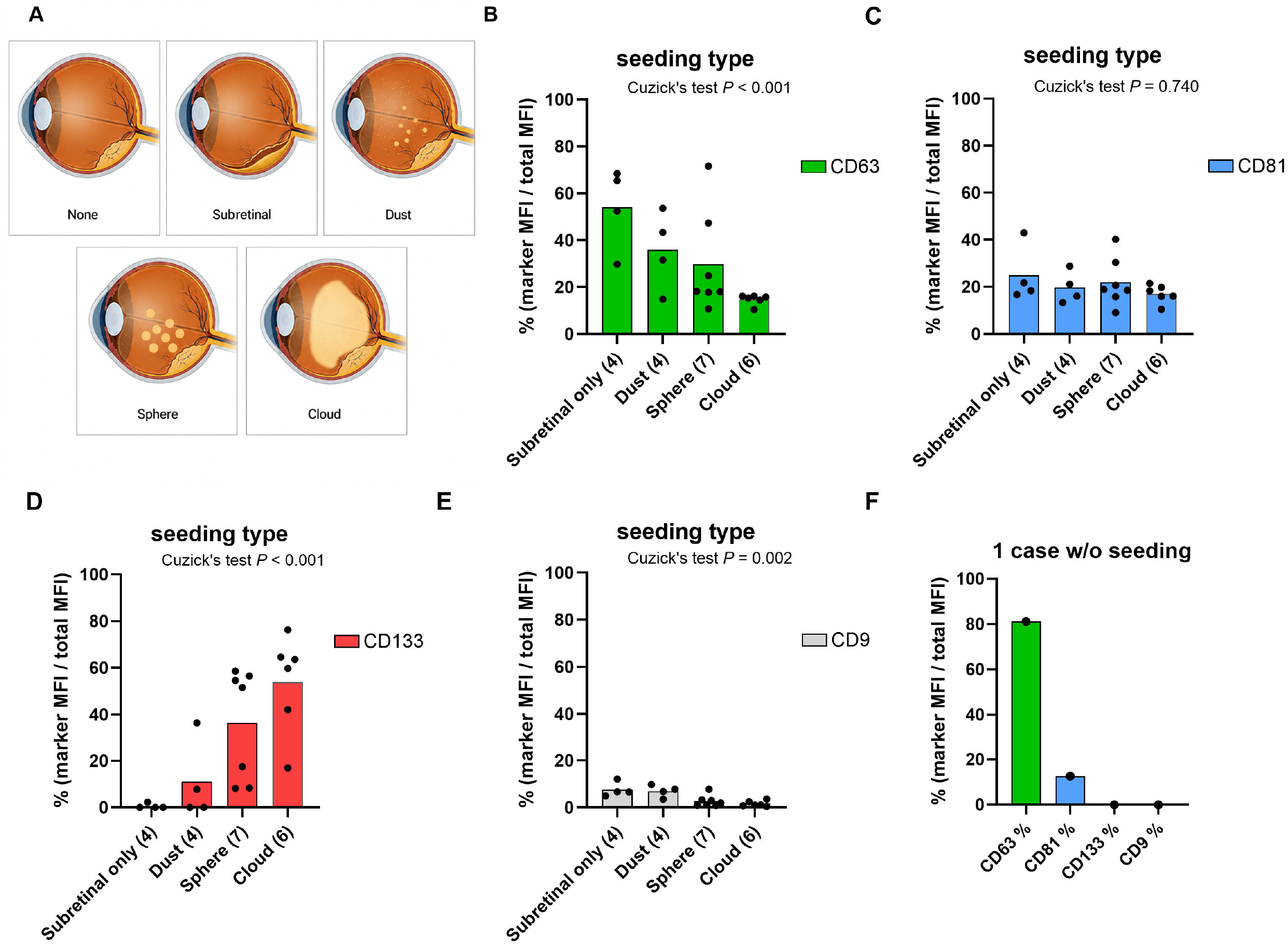
Tetraspanins and CD133 expression percentages in AH-derived EVs stratified by seeding types. (A) An illustration of different vitreous seeding types (none, subretinal only, dust, sphere, and cloud). (B-E) % MFI (marker MFI /total MFI) of CD63 (green, B), CD81 (blue, C), CD133 (red, D), and CD9 (grey, E) were measured across eyes exhibiting distinct seeding types. Each bar represents the mean ± SEM, and individual dots denote independent samples. Cuzick’s trend test was applied to assess monotonic associations across ordered seeding categories. (F) % MFI for CD63, CD81, CD133, and CD9 for included sample without seeding present.

Eyes with sphere or cloud seeding exhibited decreased EV expression of CD63 compared to eyes with no seeding or less complex types (subretinal only or dust) (Figure 4B). In contrast, CD81 expression was not affected (Figure 4C). CD133, however, showed an increasing trend associated with seeding severity (Figure 4D). CD9 also exhibited a significant decreasing trend, similar to CD63, although the proportion of the CD9 subpopulation was low (Figure 4E). Cuzick’s trend tests revealed statistically significant differences in expression for CD63 (P < 0.001), CD133 (P < 0.001) and CD9 (P = 0.002) across the four groups (Figure 4B, 4D and 4E). For the case without seeding present, the MFI percentages for CD63, CD81, CD133, and CD9 were included (Figure 4F).

To further assess the relationship between biomarker expression and seeding, we categorized samples by the seeding type and plotted MFI for CD63, CD81, CD133, CD9, and others to show signal intensities for each seeding type (Supplementary Figure 1). Although all markers exhibited progressively elevated levels with increasing seeding complexity, percentage normalization revealed the substantial contribution of distinct EV subpopulations (Figure 4). Collectively, these findings indicate that elevated EV marker expression and percentage, particularly for CD133, is most pronounced in eyes with advanced vitreous seeding patterns (sphere and cloud), supporting potential as liquid biopsy indicators of intraocular tumor burden and severity of dissemination.

### CD133^high^ EVs define a distinct tetraspanin association pattern

To investigate the relationship between CD133 expression and EV tetraspanin profiles, samples were stratified by increasing CD133 MFI, revealing heterogeneity in tetraspanin composition (Figure 5A). Notably, samples with higher CD133 MFI also demonstrated increased proportions of CD133%, while CD63%, CD81% and CD9% signals remained relatively low and stable across the cohort (Figure 5A). Based on a threshold of 10,000 MFI, samples were separated into CD133^low^ (n = 13) and CD133^high^ (n = 9) groups.

**Figure 5.**
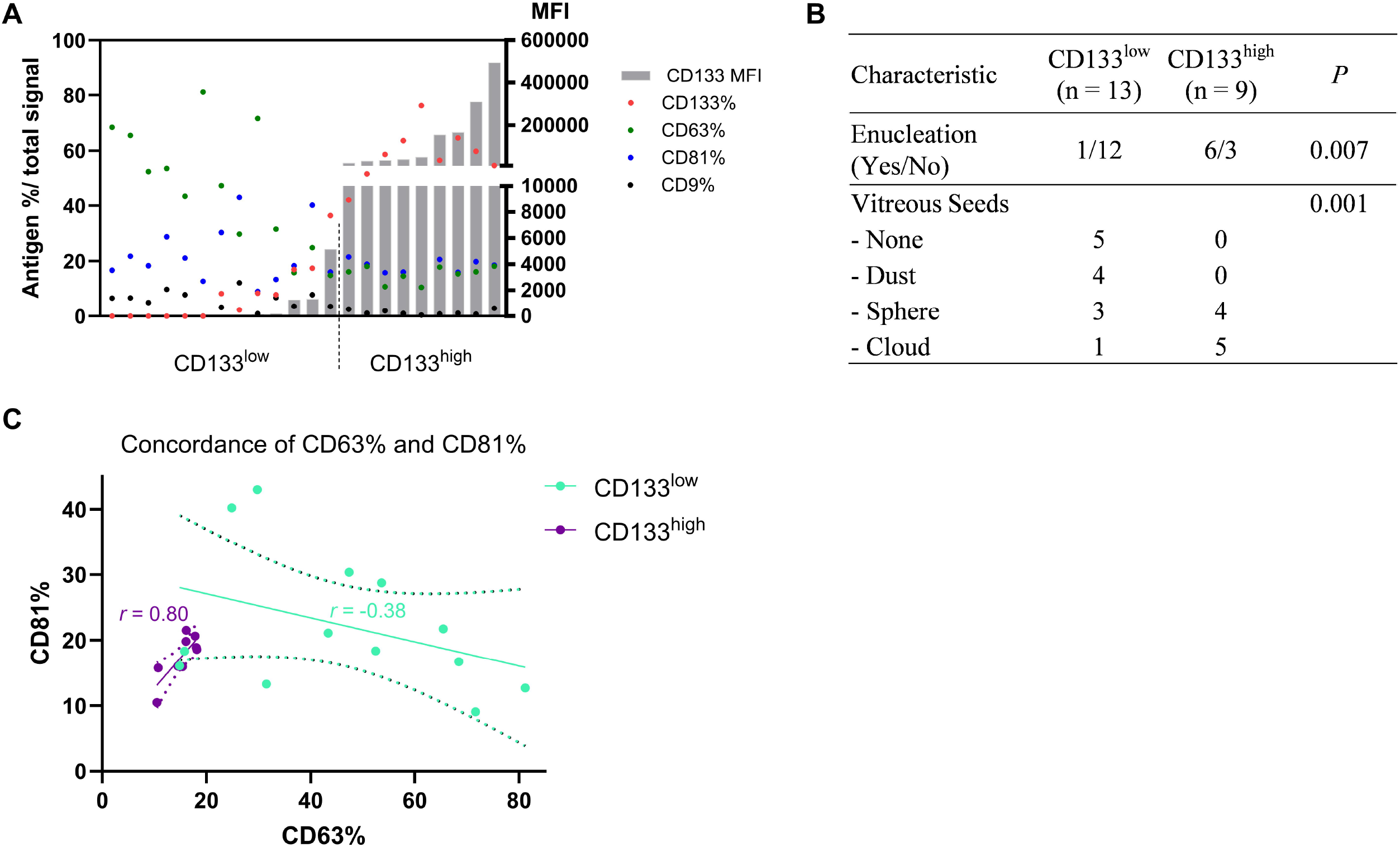
CD133 expression level defines distinct tetraspanin profiles and clinical associations in AH EVs from RB patients. (A) CD133 MFI and associated tetraspanin marker expression (% total signal) across 22 samples (gray bars, left y-axis). Dot plots overlay CD133% (red), CD63% (green), CD81% (blue), and CD9% (black) relative signal proportions (right y-axis). (B) Correlation of CD133 expression with enucleation status and vitreous seed type. Cases were separated by CD133 MFI (threshold: 10,000) into CD133^low^ (13) and CD133^high^ (9) groups. (C) A scatter plot comparing CD63% and CD81% expression across CD133^low^ (green) and CD133^high^ (purple) groups. Linear regression lines with 95% confidence intervals are shown, and Pearson’s correlation coefficient of CD63-CD81 concordances is indicated.

Clinical correlation analysis revealed a significant association between high CD133 expression and enucleation, which encompassed both primary and secondary enucleation (P = 0.007) (Figure 5B). Furthermore, CD133^high^ cases displayed a greater prevalence of aggressive vitreous seed types (P = 0.001), suggesting a potential link between elevated CD133 expression and advanced intraocular disease.

Tetraspanin co-expression patterns differed between the two groups. A scatter plot comparing CD63% and CD81% expression across CD133^low^ and CD133^high^ groups revealed strong CD63/CD81 co-expression in the CD133^high^ group (r = 0.80) that was not seen in the CD133^low^ group (r = -0.38), indicating a shift in tetraspanin co-localization patterns with increasing CD133 expression (Figure 5C).

Collectively, these findings indicate that CD133^high^ AH-derived EVs represent a phenotypically distinct subpopulation characterized by coordinated CD63 and CD81 expression, and are associated with clinically aggressive disease features, including increased enucleation rates and complex vitreous seeding patterns.

## Discussion

In this study, we investigated the association between CD133 expression on AH-derived EVs and key clinical features of RB. Analysis of a diagnostic AH cohort revealed substantial heterogeneity in CD133 signal intensity, ranging from undetectable to markedly elevated levels in advanced RB seeding phenotype. CD133 is a well-recognized biomarker for tumor-derived EVs in several malignancies, with extensive evidence in colorectal cancer linking CD133L EVs to modulation of the tumor microenvironment, promotion of tumor progression, and poorer clinical outcomes^27–30^.

Previous *in vitro* studies have highlighted the biological significance of CD133 in RB cell lines^31^, particularly its associations with stemness, proliferation, differentiation, and chemoresistance of cancer stem cells (CSC). However, the presence of CD133 on AH-derived EVs in RB and its clinical relevance have remained unclear. Our findings address this knowledge gap by correlating CD133L EV profiles with clinically relevant RB phenotypes, including seeding pattern and morphology, along with enucleation status.

Our previous work^19^ identified robust CD133 positivity in multiple RB AH-derived EV samples and RB cell lines, prompting further investigation into its origin and significance. We demonstrated that CD133 expression was consistently associated with CD63 and CD81 co-expression, a finding aligned with our prior ExoView-based analyses that established CD63/CD81 co-dominance as a marker of tumor-derived EVs in RB^17^. In the present study, this pattern was supported by observation that CD133^high^ samples demonstrated a strong CD63/CD81 co-expression signature (Figure 5C), consistent with a coordinated EV phenotype.

Notably, higher CD133 trended with more advanced IIRC stage and a greater likelihood of primary enucleation (Figure 5B), suggesting that CD133 elevation may accompany more clinically severe disease, though not functioning as an outcome predictor in this dataset.

When we looked more closely at tumor seeding characteristics, we observed differences in CD133 signal depending on the location of seeding. Eyes with vitreous seeds had significantly higher CD133 expression than those with subretinal seeding (Figure 3D). This may reflect differences in EV trafficking within the eye. The subretinal space is anatomically isolated and protected by the blood-retinal barrier, which may restrict the release or movement of EVs. In contrast, the vitreous cavity is more open and better connected to the anterior chamber, making it more likely that EVs from this region reach the AH, where we collected our samples.

We also saw that CD133^high^ eyes tended to have more sphere and cloud seeding (Figure 5B). Previous studies have demonstrated that there are higher local treatment failure risks with diffuse seeding in comparison to focal seeding^32^. Vitreous seeding is delineated into 3 distinct patterns based on the spread of tumor cells: dust, sphere, and cloud. Dust seeding is described as the result of cellular infiltration and consists of individual tumor cells, and cloud seeding is a result of translocation of the primary tumor content with a huge mass of tumor transference^20^. Sphere is described as clonal expansion from either dust or cloud but can also result directly from sprouting of the retinal tumor, forming tight clusters. Between the three classifications, differences have previously been observed related to treatment response and outcome. Dust seeding has been shown to regress more rapidly, and eyes with cloud seeding have been shown to require more injections and a higher dose than dust^33^. One study hypothesized that spheres represent an aggressive seed subtype due to the multiple layers of viable tumor cells, while cloud seeding patterns can have a lack of response to treatment due to an absence of replicating tumor cells^23^. In our cohort, CD133^+^ samples were more likely to display cloud or sphere patterns (Figure 5B), supporting a potential association between elevated CD133-EV signal and more aggressive seeding morphology.

It is important to note that multiple markers demonstrated progressively elevated total MFI levels as seeding complexity increased (Supplementary Figure 1). However, by analyzing samples according to % MFI of that specific marker over the total MFI, we were able to reveal the dominance of certain EV populations with certain seeding morphologies, paying particular attention to CD63 signal percentage in less aggressive seeding morphologies (Figure 4B) and CD133 percentage increase with the more complex seeding types (Figure 4D). While CD9 also showed a statistically significant trend with seeding morphology (Figure 4E), its % MFI was low in comparison to other markers measured. Notably, samples with higher CD133 MFI also demonstrated increased proportions of CD133%, while CD63%, CD81% and CD9% signals remained relatively low and stable across the cohort (Figure 5A).

Collectively, these findings indicate that both elevated EV marker expression and percentage, particularly for CD133, are most pronounced in eyes with advanced vitreous seeding patterns (sphere and cloud), supporting their potential as liquid biopsy indicators of intraocular tumor burden and severity of dissemination.

Although CD133 percentage rose in parallel with absolute CD133 intensity, suggesting a consistent biological signal, our findings do not establish CD133 as a mechanistic driver of RB dissemination. Rather, CD133 likely marks an EV subpopulation enriched in eyes with more extensive or advanced seeding.

The ability to detect aggressive seeding patterns through a minimally invasive AH tap could alter clinical decision making. Eyes with high CD133□ EV levels might warrant more intensive local therapy, earlier intravitreal chemotherapy, or closer follow-up intervals, even if traditional imaging findings are equivocal. Such an approach could help preempt treatment failure and reduce enucleation rates. However, prospective validation, ideally with longitudinal sampling, is required before any clinical application.

This study has several limitations, including a modest, single-institution sample size and the absence of direct validation of EV origin. While CD133□ EVs were interpreted as tumor-associated based on co-expression with canonical markers and clinical correlations, further functional studies are needed to confirm this. The use of a median-based CD133 cutoff was exploratory and required refinement in larger cohorts. Additionally, anatomical differences in EV access to the AH may bias detectability toward vitreous rather than subretinal disease. Variability in EV abundance across clinical samples may also influence MFI values, although percentage normalization partially mitigates this.

Despite these limitations, our findings provide novel evidence linking AH-derived EVs to clinically relevant tumor seeding features and enucleation status in RB. These findings, along with the concordance between CD133 and CD63/81 signal, suggest that CD133^+^ EVs may represent a specific subpopulation tied to tumor biology that carries useful clinical information and thus could be explored further as non-invasive progression biomarkers. With further validation and mechanistic insight, CD133□ EVs hold promise as markers to support risk stratification and guide treatment planning in clinical practice.

## Acknowledgements

The authors would like to acknowledge the Flow Cytometry Core of the Saban Research Institute at Children’s Hospital Los Angeles for their contributions to this research. We would also like to thank Brianne Brown, Drishti Pandya and Hope Galarneau for their hard work.

## Supplementary Material

**Supplementary Figure S1. Tetraspanins and CD133 expression levels in AH-derived EVs stratified by vitreous seeding types**. MFI of CD63 (green), CD81 (blue), CD133 (red), CD9 (grey) and others (dark grey) were measured for all samples, which were grouped by distinct seeding types (no seeding, subretinal, dust, sphere, cloud). Cuzick’s non-parametric trend test to assess monotonic changes across increasing seeding complexity.

