## Supplementary figures and images for "Extracellular Vesicle Biomarkers in the Aqueous Humor Correlate with Retinoblastoma Seeding"

### Supplementary Figure 1

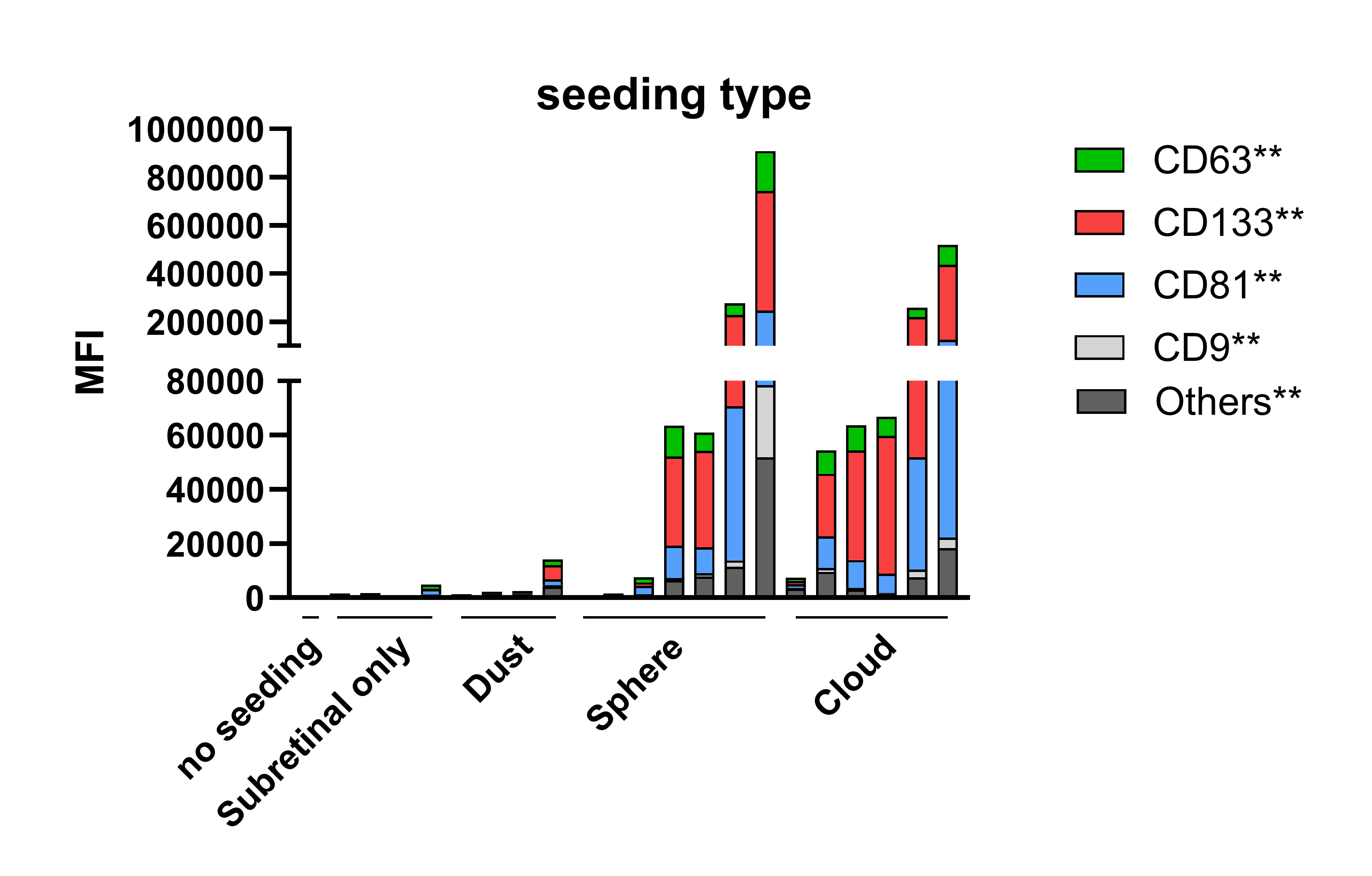
